# *Candida albicans* Isolates 529L and CHN1 Exhibit Stable Colonization of the Murine Gastrointestinal Tract

**DOI:** 10.1101/2021.06.27.450080

**Authors:** Liam McDonough, Animesh Anand Mishra, Nicholas Tosini, Pallavi Kakade, Swathi Penumutchu, Shen-Huan Liang, Corrine Maufrais, Bing Zhai, Ying Taur, Peter Belenky, Richard J. Bennett, Tobias M. Hohl, Andrew Y. Koh, Iuliana V. Ene

## Abstract

*Candida albicans* is a pathobiont that colonizes multiple niches in the body including the gastrointestinal (GI) tract, but is also responsible for both mucosal and systemic infections. Despite its prevalence as a human commensal, the murine GI tract is generally refractory to colonization with the *C. albicans* reference isolate SC5314. Here, we identify two *C. albicans* isolates, 529L and CHN1, that stably colonize the murine GI tract in three different animal facilities under conditions where SC5314 is lost from this niche. Analysis of the bacterial microbiota did not show notable differences between mice colonized with the three *C. albicans* strains. We compared the genotypes and phenotypes of these three strains and identified thousands of SNPs and multiple phenotypic differences, including their ability to grow and filament in response to nutritional cues. Despite striking filamentation differences under laboratory conditions, however, analysis of cell morphology in the GI tract revealed that the three isolates exhibited similar filamentation properties in this *in vivo* niche. Notably, we found that SC5314 is more sensitive to the antimicrobial peptide CRAMP, and the use of CRAMP-deficient mice increased the ability of SC5314 to colonize the GI tract relative to CHN1 and 529L. These studies provide new insights into how strain-specific differences impact *C. albicans* traits in the host and advance CHN1 and 529L as relevant strains to study *C. albicans* pathobiology in its natural host niche.

**IMPORTANCE:** Understanding how fungi colonize the GI tract is increasingly recognized as highly relevant to human health. The animal models used to study *Candida albicans* commensalism commonly rely on altering the host microbiome (via antibiotic treatment or defined diets) to establish successful GI colonization by the *C. albicans* reference isolate SC5314. Here, we characterize two *C. albicans* isolates that can colonize the murine GI tract without antibiotic treatment and can therefore be used as tools for studying fungal commensalism. Importantly, experiments were replicated in three different animal facilities and utilized three different mouse strains. Differential colonization between fungal isolates was not associated with alterations in the bacterial microbiome but rather with distinct responses to CRAMP, a host antimicrobial peptide. This work emphasizes the importance of *C. albicans* intra-species variation as well as host anti-microbial defense mechanisms in defining commensal interactions.

## INTRODUCTION

The fungal component of the human microbiota, the mycobiota, is increasingly recognized as playing key roles in host homeostasis (1–7). *Candida albicans*, a pathobiont that is found in over 70% of individuals, is a prominent member of the gastrointestinal (GI) mycobiota (8, 9). This species is present in multiple niches of the human body and can cause a variety of opportunistic mucosal and systemic infections. Disseminated infections can arise when *Candida* cells in the GI tract translocate into the bloodstream (10, 11), as has been observed in murine models of mucositis and neutropenia (12) and in patients undergoing allogeneic hematopoietic cell transplants (13). *C. albicans*, as well as other *Candida* species, are also linked to intestinal disease, with *C. albicans* consistently found at high levels in cohorts of Crohn’s Disease and ulcerative colitis patients (14). The loss of host signaling pathways involved in fungal recognition, such as those involving Dectin-1 or Dectin-3, may exacerbate colitis due to increased *Candida* levels in the gut (15, 16).

The impact of the GI mycobiota is not limited to gut mucosal tissues but can also modulate systemic responses distal to this organ. For example, *C. albicans* cells in the GI tract can drive the induction of systemic Th17 responses in both mice and humans (1, 4). These systemic responses can be a double-edged sword as they can provide protection against systemic infections by fungi or other microbial pathogens but can cause increased airway inflammation in response to antigens in the lung (1, 4). Understanding of GI colonization by *C. albicans* and related fungal species therefore has far reaching consequences for understanding immune homeostasis at both intestinal sites and sites distal to the gut.

Given their central role in host homeostasis, it is notable that most laboratory mice are not readily colonized with *C. albicans* or other fungi (17, 18). The importance of commensal fungi to the biology of laboratory mice was highlighted in a recent study in which lab mice were rewilded by release and subsequent recapture from an outdoor facility (17). Notably, rewilded mice showed enhanced differentiation of memory T cells and increased levels of circulating granulocytes, and these changes were associated with increased fungal colonization of the GI tract (17). Inoculation of lab mice with fungi from rewilded mice or with *C. albicans* was sufficient to enhance immune responses, further establishing that the gut mycobiota can play broad roles in educating host immunity.

Relatively little is known about the fungal and host mechanisms that regulate GI colonization by species such as *C. albicans*. Most studies have relied on antibiotic supplementation to allow the standard ‘laboratory’ strain of *C. albicans*, SC5314, to stably colonize the GI tract of mice (12, 19, 20). Several other *Candida* strains are also unable to colonize the murine GI tract without the use of antibiotics, including *C. albicans* strains WO-1, Can098, 3153A, ATCC 18804, OH-1, *Candida glabrata* ATCC 15126, a *Candida parapsilosis* clinical isolate, and *Candida tropicalis* ATCC 66029 (21–23).

Antibiotic treatment against bacterial taxa can enable fungal colonization as specific bacterial commensals induce the transcription factor HIF-1α in enterocytes which in turn leads to production of CRAMP, an antimicrobial peptide related to the human cathelicidin LL-37 (21). LL-37 has been shown to exhibit both antibacterial and antifungal activity (24), can inhibit *Candida* adhesion and affect cell wall integrity by interacting with cell wall components, including the exoglucanase Xog1 (25–27). CRAMP kills *C. albicans* cells *in vitro* (28) and inhibits GI colonization, as shown by increased *C. albicans* colonization in mice lacking CRAMP (21). Conversely, on the fungal side, loss of filamentation has been linked to enhanced GI colonization by *C. albicans* cells in both antibiotic-treated and germ-free mice (29–32). Several transcriptional regulators of the *C. albicans* mating circuit have also been shown to impact fungal fitness levels in this niche (31, 33–35).

While SC5314 represents the standard isolate of *C. albicans* used by many in the field, several studies have established that *C. albicans* isolates display a wide range of phenotypic properties both *in vitro* and in models of infection (36–40). Intra-species variation can therefore have a major impact on *C. albicans* strain behavior and determine the outcome of host-fungal interactions. Understanding inter-strain differences is critical for determining the breadth of properties displayed by a species and could lead to new insights into mechanisms of fungal adaptation, niche specificity and pathogenesis (33, 41–44).

Here, we compared the ability of different *C. albicans* strains to colonize the murine GI tract without antibiotic treatment. We identified two isolates, 529L and CHN1, that stably colonize the GI tract under conditions where SC5314 is consistently lost from this niche. Similar findings were obtained when using three different mouse lines in three different animal facilities, highlighting the robustness of this finding. 529L and CHN1 also outcompeted SC5314 in direct competition experiments in the murine intestine, establishing that these strains exhibit an increased relative fitness for this niche. Analysis of the phenotypic properties of SC5314, CHN1, and 529L revealed stark differences in filamentation and metabolism between these strains *in vitro*. However, filamentation differences were not evident in the murine gut, highlighting how *in vivo* phenotypes can differ from those observed *in vitro*. Instead, we show that CHN1 and 529L were more resistant to killing by the CRAMP peptide relative to SC5314 and linked these differences to GI colonization fitness using mice lacking CRAMP. Together, these studies highlight how differences between *C. albicans* isolates can dictate differences in gut colonization and establish CHN1 and 529L as relevant tools for the study of this fungus in its commensal niche.

## RESULTS

### *C. albicans* strains 529L and CHN1 can stably colonize the murine GI tract without antibiotics

We compared the ability of three *C. albicans* human isolates to each colonize the GI tract of three different strains of mice in the absence of antibiotic supplementation. The isolates tested were SC5314, the standard ‘laboratory’ isolate originally obtained from a bloodstream infection (45), 529L, isolated from the oral cavity (46), and CHN1, isolated from the lung (47). These strains were orally gavaged to BALB/c (Charles River Laboratories), C57BL/6J (Jackson Laboratories) and C3H/HeN (Envigo), mice fed a standard chow diet in animal facilities in Texas (TX), New York (NY), or Rhode Island (RI). GI colonization levels were monitored by plating mouse fecal pellets every 2-7 days.

SC5314 did not stably colonize the GI of any of the mice tested. For example, CFU levels decreased 1-2 logs in the first 7-14 days of infection in both C3H/HeN (TX) and C57BL/6J (RI) mice and fell below detection levels at later time points (Figure 1A-D). In contrast, 529L and CHN1 more stably colonized the GI tract in each of the mouse strain backgrounds, particularly in C57BL/6J (RI, NY) and C3H/HeN (TX) mice, being maintained for 28-48 days post inoculation (Figure 1A-D). For C57BL/6J (RI, NY) mice, colonization differences between isolates were readily apparent in the first week post gavage and in the RI facility these differences increased out to 28 days of colonization (Figure 1A, B). Most C67BL/6J mice cleared SC5314 cells whereas CHN1 and 529L were present at >10^3^CFUs/g feces in both the TX and RI facilities at the end of the experiment (Figure 1A-B and Supplemental Figure 1A). In BALB/c (RI) mice, 529L and CHN1 had higher levels of colonization than SC5314 during the first 5 days, although significance was not observed at later time points (Figure 1C). Finally, in C3H/HeN (TX) mice, both 529L and CHN1 were present at higher levels than SC5314 throughout the time course with SC5314 cells no longer recovered from the fecal pellets of any mice by day 21 (Figure 1D and Supplemental Figure 1A).

**Figure 1.**
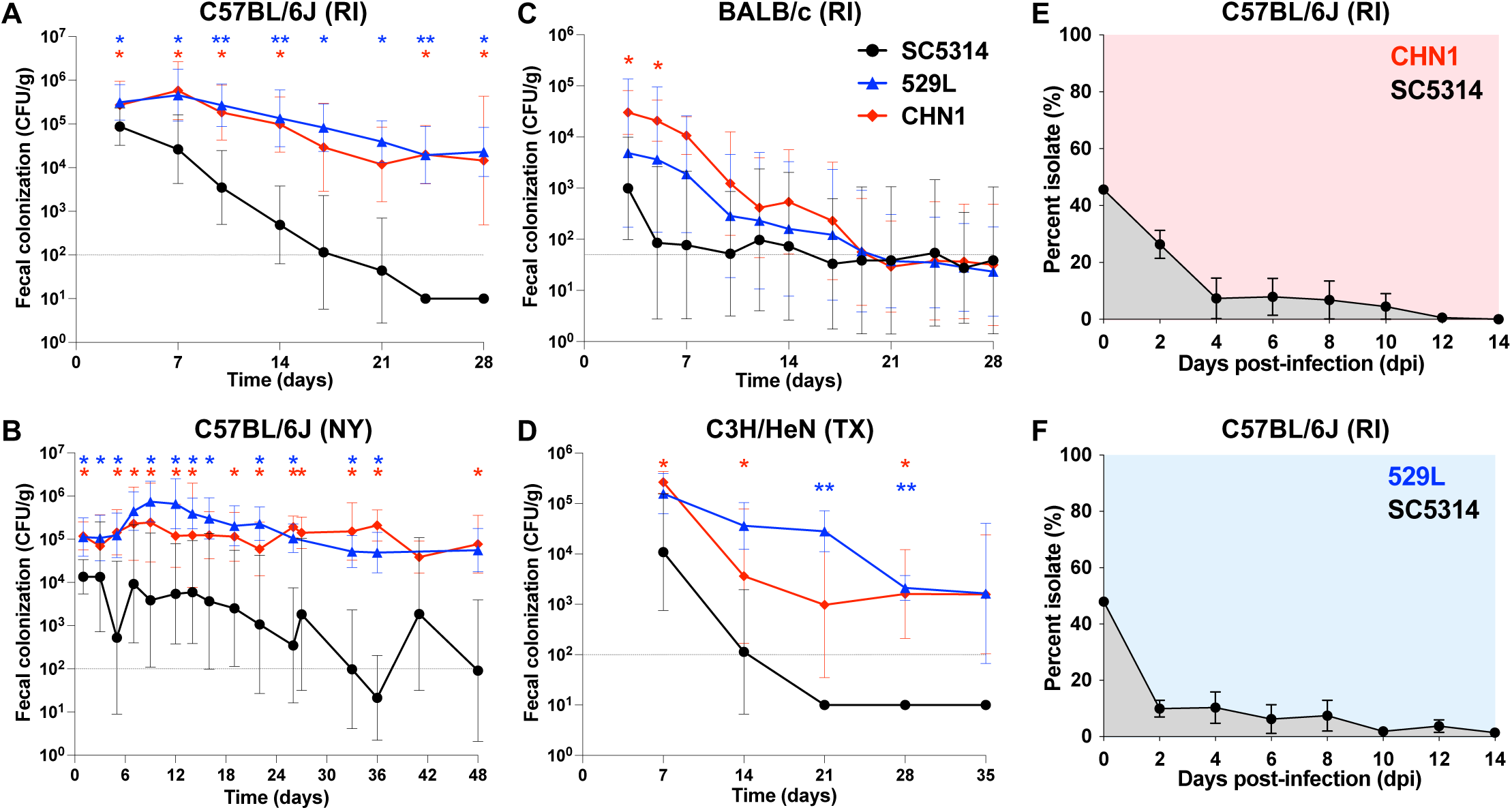
*C. albicans* isolates 529L and CHN1 can stably colonize the gastrointestinal tract of C57BL/6J (A - RI, n = 8 mice; B - NY, n = 10-18 mice), BALB/c (C - RI, n = 8 mice) and C3H/HeN (D - TX, n = 8 mice) mice without antibiotic treatment. Panels show geometric means with 95% CI of fecal colonization levels (CFUs/g) over time. Asterisks reflect comparisons between isolates at individual time points using Mann-Whitney tests; *, *P* < 0.05, **, *P* < 0.01. Dotted horizontal lines indicate the minimum CFU detection level for each experiment. (E-F) Direct competitions between SC5314 and 529L (E) or CHN1 (F) in the GI of C57BL/6J mice (RI). Isolates were co- inoculated in a 1:1 ratio and their proportions were determined using nourseothricin selection upon recovery from fecal pellets. Panels show means ± SEM from 4 single housed mice.

At the end of the experiment, colonization levels were also examined by recovery of colony forming units (CFUs) from GI organs in C57BL/6J and BALB/c mice. Analysis revealed relatively high levels of 529L and CHN1 present in C57BL/6J (RI, NY) organs, while SC5314 was typically not recovered from any GI organs (Supplemental Figure 1B). For BALB/c (RI) mice, we could not identify significant differences in organ colonization levels between the three isolates at day 28, reflecting the fact that each isolate showed reduced colonization at these time points in this mouse background (Supplemental Figure 1B).

Having established that 529L and CHN1 showed increased fitness relative to SC5314 in mono-colonization experiments, we tested whether these strains showed fitness differences in direct competition experiments. To distinguish the strains in a direct competition, SC5314 was transformed with a nourseothricin resistance gene (*SAT1*) targeted to the *NEUT5L* on chromosome (Chr) 5, which is a neutral locus for integration of ectopic constructs (48). To verify that the presence of *SAT1* at this site does not alter the fitness of isolates during gut colonization, we performed competitions between SC5314 and two independently transformed SC5314 isolates containing *SAT1*. A 1:1 mix of *SAT1*-marked and unmarked SC5314 was introduced into the GI and relative strain abundance determined by calculating the proportion of nourseothricin-resistant colonies recovered from fecal pellets over 14 days. Experiments revealed no significant advantage between the two versions of SC5314, indicating that the presence of *SAT1* did not affect *C. albicans* fitness in the GI tract (Supplemental Figure 2A-B).

Next, a 1:1 mix of SC5314 (*SAT1*-marked) and 529L or CHN1 was introduced into the GI tract and the relative proportions of each strain determined from fecal pellets of C57BL/6J (RI) mice. By day 4 post gavage, both 529L and CHN1 began to dominate the colonizing population, with SC5314 cells representing less than 10% of CFUs in fecal pellets (Figure 1E-F). By day 14, CHN1 and 529L accounted for 100% and 98.5% of the cells recovered from the feces, respectively, and similar proportions of isolates were observed across the different GI organs (stomach, small intestine, colon, and cecum; Supplemental Figure 3A-B). These results indicate that both CHN1 and 529L display increased competitive fitness relative to SC5314 throughout the GI tract.

To extend these findings, we performed similar competition experiments using both *SAT1*-marked and unmarked versions of each isolate in different combinations in the C57BL/6J (NY) murine background. Experiments confirmed the RI facility findings in that both CHN1 and 529L showed increased competitive fitness relative to SC5314 (Supplemental Figure 4A-B). Experiments also showed that CHN1 exhibited a consistent fitness advantage over 529L (Supplemental Figure 4C). We noted, however, that large fluctuations were observed in the overall proportions of each isolate in these competitions. When competing CHN1 and SC5314, differences between strains were apparent approximately 24 days post gavage, when CHN1 became dominant in fecal pellets (Supplemental Figure 4A). For 529L versus SC5314 competitions, 529L represented ∼80% of the fungal population after 3 days or after 22 days depending on which strain carried the *SAT1* gene (Supplemental Figure 4B). Similarly, the fitness advantage of CHN1 relative to 529L was evident much earlier in one competition than in the other (Supplemental Figure 4C), illustrating variability in the dynamics of gut colonization. Despite this variation, these findings establish that CHN1 and 529L consistently show increased fitness in the murine GI tract compared to SC5314.

### *C. albicans* isolates do not significantly affect the composition of the gut bacterial microbiome

Commensal bacterial gut microbiota (particularly the phylum Bacteroidetes and family Lachnospiraceae) are important for murine resistance to *C. albicans* colonization (21), an association that was recently corroborated in adult hematopoietic cell transplant recipients (13). To assess the impact of colonization with different *C. albicans* isolates on the bacterial microbiota, the 16S rRNA hypervariable region was sequenced from fecal pellets from both BALB/c and C57BL/6J mice colonized with these isolates and was performed in mice housed in different animal facilities (RI and NY). We found that colonization with SC5314, 529L, or CHN1 strains did not result in any significant differences in microbiome composition at phylum or family levels in either animal facility (Figure 2A-C). Colonization also did not significantly affect the alpha diversity of the bacterial microbiome as measured by the Shannon Diversity Index or cause significant changes in beta diversity, with samples displaying no significant clustering on the PCoA projection of Bray-Curtis distances for both BALB/c (Supplemental Figure 5A) and C57BL/6J mice (Supplemental Figure 5B-C) in the two facilities (RI, NY). These experiments establish that *C. albicans* colonization with these different isolates has a minimal impact on the composition of the bacterial microbiota under the conditions evaluated in this study.

**Figure 2.**
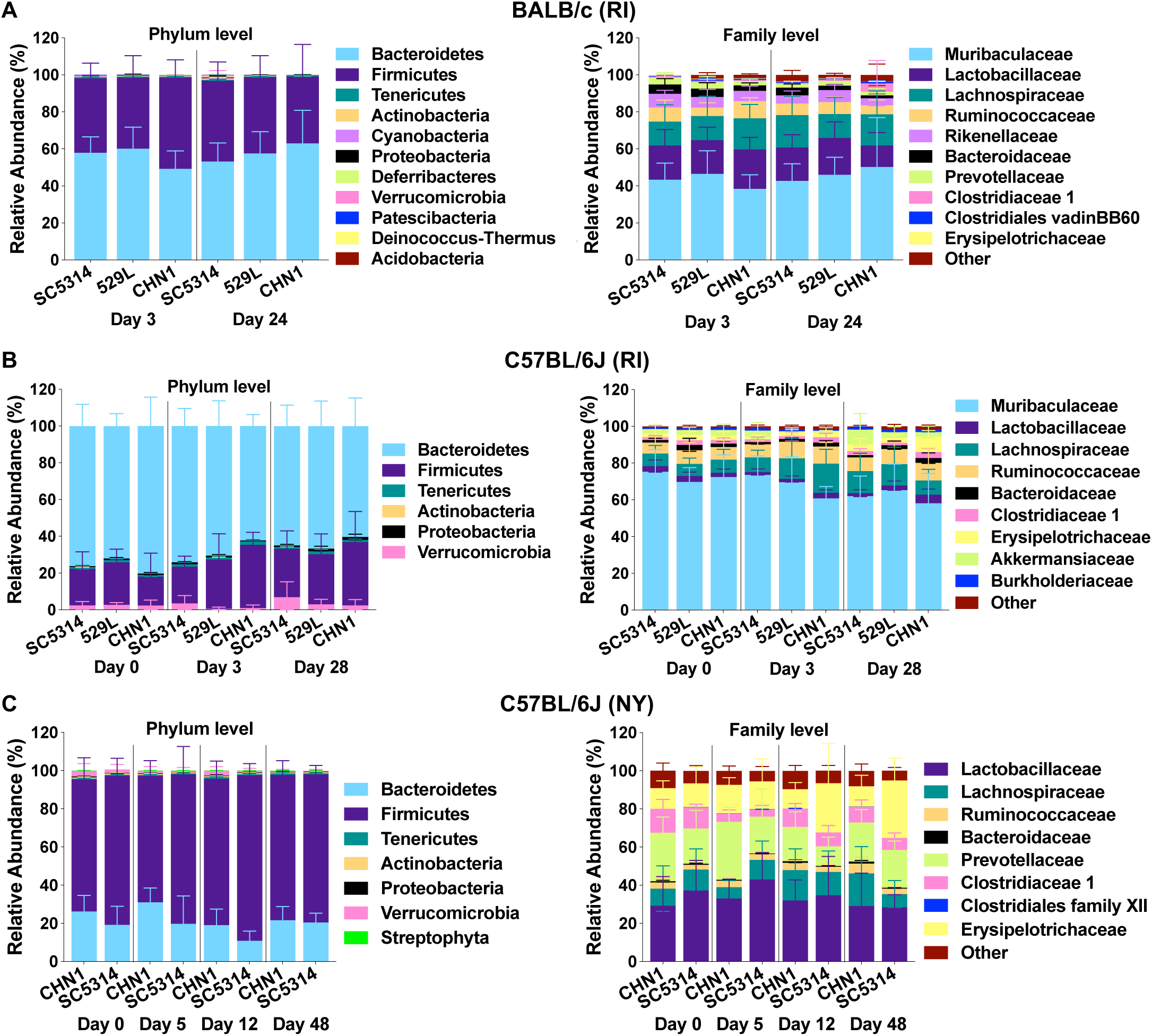
Microbiome composition of BALB/c (A, RI) and C57BL/6J (B, RI; C, NY) mice colonized with *C. albicans* isolates SC5314, CHN1 and 529L. Plots show microbiome relative abundances at the phylum and family levels for mice from Figure 1. Day 0 time points indicate the microbiome composition prior to *Candida* gavage.

### SC5314, 529L and CHN1 show distinct metabolic and filamentation properties *in vitro*

To investigate the mechanism by which 529L and CHN1 exhibit increased GI fitness relative to SC5314 we compared the phenotypes of the three isolates in a series of *in vitro* assays. First, colony filamentation was examined on YPD, SCD, Spider and Lee’s + glucose media (Figure 3A). Cells were plated on these media, incubated at 37°C for 4 days, and assessed for filamentation. No visible differences were noted between the three strains when grown on YPD, however 529L displayed markedly reduced colony filamentation on SCD, Spider and Lee’s + glucose medium relative to both SC5314 and CHN1. Next, cell morphology was examined in various liquid filamentation-inducing conditions after 6 h of growth at 37°C. Consistent with colony phenotypes, 529L did not efficiently filament under these conditions and only formed rare hyphae or pseudohyphae in medium supplemented with 10% fetal calf serum (Figure 3B). In contrast, SC5314 and CHN1 displayed a strong filamentation response across all media tested.

**Figure 3.**
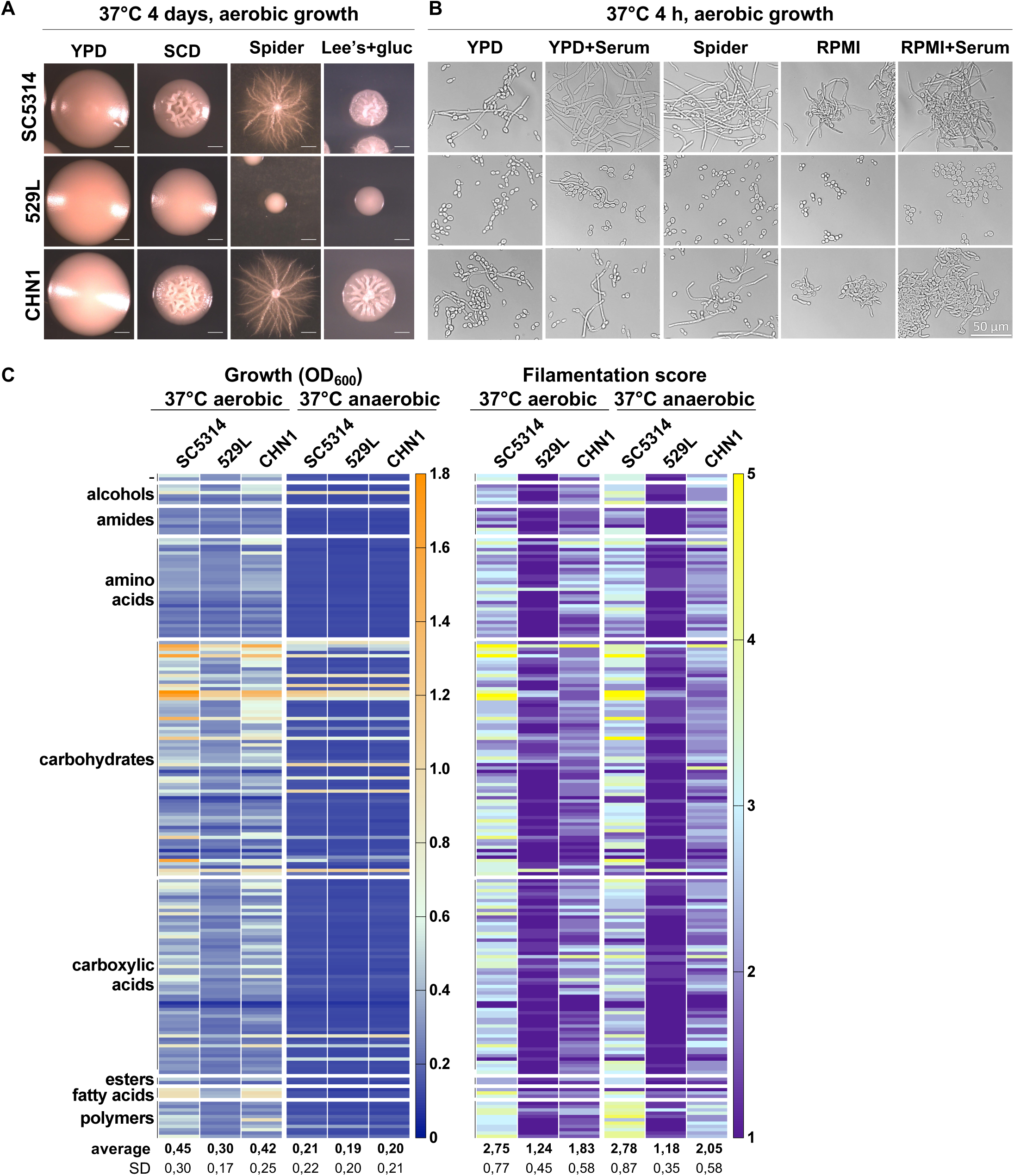
*In vitro* growth and filamentation of isolates SC5314, CHN1, 529L in different laboratory media and nutritional conditions. Colony (A) and cell (B) morphology of isolates grown at 37°C under aerobic conditions in different laboratory media. Scale bars, 1 mm (A) and 50 μm (B). (C) Growth and filamentation of isolates SC5314, CHN1, 529L on Biolog carbon source plates (PM01-02) under aerobic and anaerobic conditions. Carbon sources are grouped according to their biochemical group. After 24 h of growth at 37°C, each well was scored for filamentation on a 1 to 5 scale (1 and 5 represent conditions where 0-20% and 80-100% of cells showed visible filamentation, respectively). Bottom tables indicate means ± SD from two biological replicates for each condition.

To further characterize phenotypic differences between these strains, we utilized Phenotype Microarrays (PM, Biolog) containing a set of 190 different carbon sources. The three isolates were seeded on PM plates and incubated with shaking at 37°C for 24 h, both aerobically and anaerobically. Growth was evaluated by taking biomass (OD_600_) readings and individual wells were assessed for filamentation using a semi-quantitative score of 1 to 5. A score of 1 indicates 0-20% hyphae observed, while a score of 5 indicates that 80-100% of the population formed hyphae. Under aerobic conditions, the three isolates displayed different growth capacity across carbon sources, with SC5314 being able to reach a higher biomass (mean OD_600_ across wells = 0.45 ± 0.3 SD) than both 529L (0.30 ± 0.17) and CHN1 (0.42 ± 0.25) (Figure 3C, *P* < 0.01 for both isolates relative to SC5314, df = 191, two-way Anova). Although smaller, differences in growth were also apparent when the isolates were incubated anaerobically (Figure 3C, *P* < 0.0001 for both isolates). Analysis of filamentation under aerobic conditions revealed that 529L again displayed a severe filamentation defect across the surveyed carbon sources (average filamentation score 1.2 ± 0.5), while CHN1 displayed an intermediate filamentation capacity (1.8 ± 0.6) compared to SC5314 (2.8 ± 0.8, Figure 3C, *P* < 0.0001 for both isolates). Similar results were observed when comparing filamentation of the three isolates under low oxygen conditions (Figure 3C, *P* < 0.0001 for both isolates).

Increased *C. albicans* GI colonization has been previously associated with decreased levels of short chain or medium chain fatty acids (49, 50). In addition, the presence of short chain carboxylic acids has been shown to reduce *C. albicans* filamentation by modulating external pH and this effect could promote gut colonization (51–54). We therefore assessed the impact of carboxylic acids (acetic, butyric, lactic, capric, succinic, propionic, and citric acid) on the ability of the three strains to grow and form filaments. Aerobic growth on short chain carboxylic acids revealed that CHN1 and 529L showed reduced filamentation relative to SC5314, with a larger defect observed for 529L (*P* < 0.01 for both isolates relative to SC5314, df = 9, two-way Anova), which also displayed reduced growth (*P* < 0.05, Supplemental Figure 6A). Differences in filamentation were also observed for 529L under anaerobic conditions (*P* < 0.01), where isolates showed similar growth levels on this subset of carboxylic acids (Supplemental Figure 6A).

While it is possible that reduced filamentation could simply be the result of reduced growth, a correlation analysis between growth and filamentation levels across all carbon sources tested revealed that this was not the case (Supplemental Figure 6C). This was most apparent when the three isolates were grown under anaerobic conditions - a simple linear regression resulted in a goodness of fit with R^2^of 0-0.14, indicating the absence of a correlation between growth and filamentation across these conditions (Supplemental Figure 6C). Overall, these results indicate that both 529L and CHN1 have reduced *in vitro* growth and filamentation capacities relative to SC5314, with these differences being more pronounced for 529L.

### SC5314, 529L and CHN1 show extensive genetic differences

Previous reports have associated the presence of *C. albicans* aneuploid chromosomes with increased fitness for particular host niches, including trisomy of Chr 6 which was repeatedly selected for during oral infection (55) and trisomy of Chr 7 which favored colonization of the mouse GI tract (56). Thus, we examined the whole genome sequences of the three isolates to identify large and small genetic changes that could contribute to differential colonization of this niche. 529L and SC5314 have been previously sequenced (37, 57), therefore only CHN1 was *de novo* sequenced for this study. All three isolates were compared to the SC5314 reference genome (assembly 22) and comparative genomic analyses were performed between CHN1/529L and the SC5314 version examined here. Phylogenetic analysis revealed that CHN1 and 529L isolates belong to the relatively rare clades A and 16, respectively, being distinct from SC5314 which belongs to clade 1 (37, 41). This analysis also reveals that 529L and CHN1 are more closely related to each other than they are to SC5314 (41).

We found that all three isolates were euploid across all chromosomes (Supplemental Figure 7A), eliminating aneuploidy of specific chromosomes as a potential explanation for differences in GI fitness. However, the isolates displayed differences in heterozygosity patterns across their genome, with large homozygous regions (01-0.87 Mbp) present on multiple chromosomes (Supplemental Figure 7B-C). Certain homozygous regions were shared in the 529L and CHN1 whole genome sequences, with telomeric regions of Chr 7R and Chr RR displaying minimal heterozygosity (Supplemental Figure 7B-C). Variant calling comparing 529L and CHN1 with SC5314 revealed 112,057 and 86,513 variants, respectively (Supplemental Figure 7D and detailed in Supplemental Tables 3-4). Approximately 48% of all variants were found in coding regions, ∼90% of the total variants were represented by SNPs while the remaining 10% represented insertions/deletions (Supplemental Figure 7D). Given the large number of genetic differences present between isolates, identification of variants associated with increased stability in the host GI tract would require extensive functional analyses which are beyond the scope of the current study.

### SC5314, 529L, and CHN1 display similar morphologies in the GI tract

Since the ability of *C. albicans* to colonize the mammalian GI tract is associated closely with its propensity to filament (29, 30, 32, 33, 35, 58, 59), we directly assessed the morphology of CHN1, 529L and SC5314 cells in the gut of C57BL/6J mice (RI). We utilized an antibiotic model of gut colonization to facilitate higher levels of fungal colonization than an antibiotic-free model thereby enabling morphotypic analysis of fungal cells in GI tissue sections (see Supplemental Figure 8 for fecal and organ fungal burdens). Consistent with previous studies (32, 35), analysis of colon tissue sections colonized with SC5314 showed the presence of both yeast and filamentous forms (Figure 4A). Colonization with 529L and CHN1 also revealed the presence of both morphological forms, both in the lumen and near the colon epithelium (Figure 4A). Quantification of yeast and filamentous cells from different segments of the GI tract revealed that 529L exhibited higher proportions of filamentous cells than SC5314 in the jejunum (18% more filamentous), but similar proportions of filamentous cells in the other GI segments (Figure 4B). This result was unexpected given that 529L was defective for filamentation under most *in vitro* growth conditions (Figure 3). In turn, CHN1 showed reduced filamentation in the duodenum relative to SC5314 (29% fewer filamentous cells) but the opposite trend in the colon (3% more filamentous cells; Figure 4B). This data demonstrates that, in general, clinical isolates 529L and CHN1 display a similar overall distribution of yeast and hyphal forms to SC5314 when colonizing the murine GI tract. The absence of consistent differences in cell morphology between the three isolates *in vivo* indicates that filamentation *per se* does not appear to drive differences in GI colonization.

**Figure 4.**
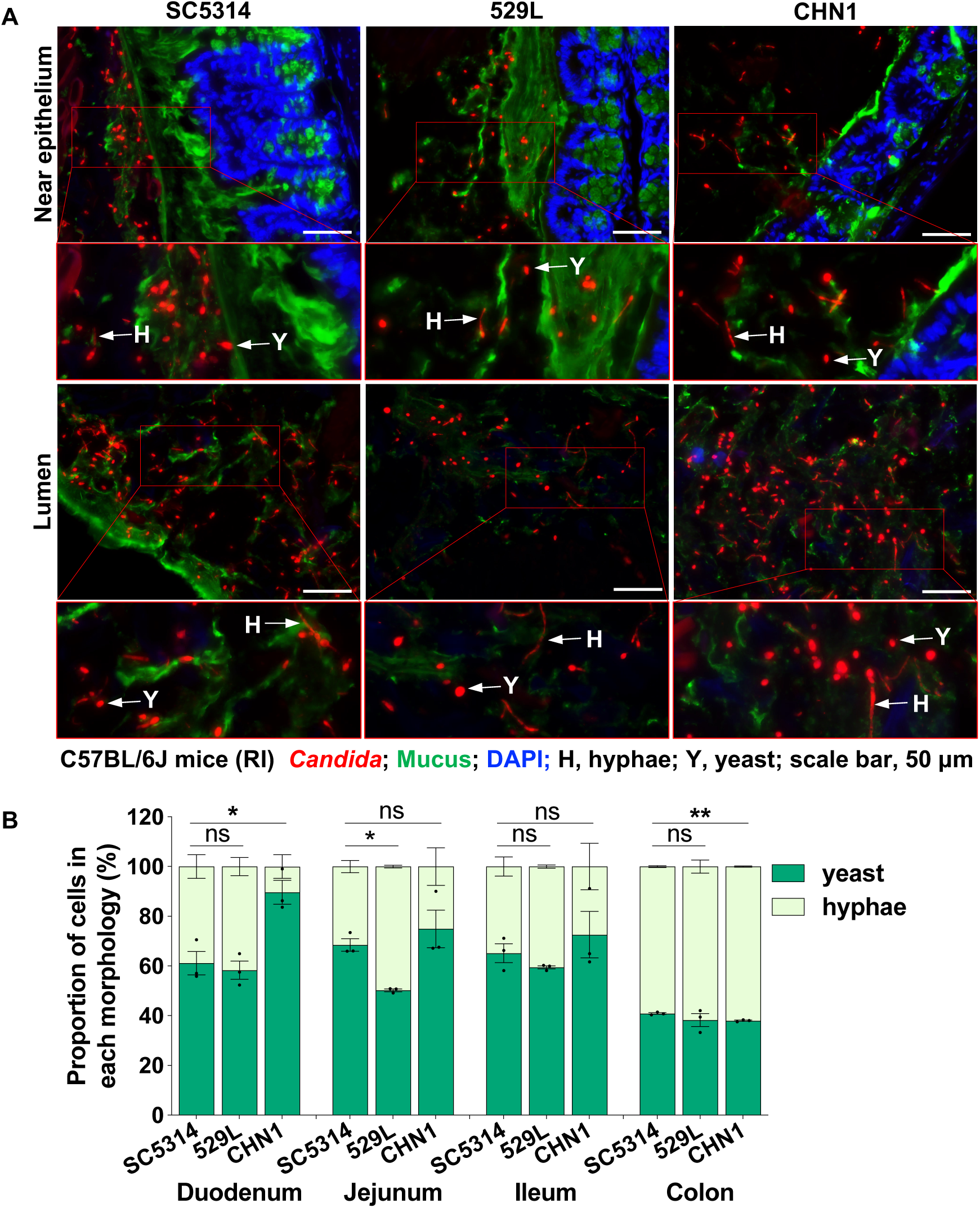
Morphology of *C. albicans* cells in the GI of C57B/6J mice (RI) using an antibiotic model of colonization (n = 3 mice, single-housed). (A) FISH-stained *Candida* cells from colon sections. The colon tissues from mice were stained with Cy3-coupled 28S rRNA fungal probe to stain both yeast and hyphal cells. Epithelium and mucus were stained with DAPI and UEA1 and WGA1 coupled with fluorescein, respectively. Scale bar, 50 μm. Arrows indicate different *Candida* cell morphologies - H, hyphae; Y, yeast. (B) *Candida* cells in the different GI sections of C57B/6J mice (RI) on antibiotics were stained with anti-*Candida* antibody coupled with FITC. Histograms show the proportion (%) of yeast and hyphal cells in different GI organs (means ± SEM). Asterisks indicate statistical significance using unpaired parametric t-tests, * *P* < 0.05, ** *P* < 0.01; ns, not significant, n = 50-600 cells per section.

### 529L and CHN1 exhibit increased resistance to the antimicrobial peptide CRAMP relative to SC5314

The intestinal epithelial-derived antimicrobial peptide CRAMP was previously shown to inhibit *C. albicans* colonization in the murine GI tract (21). We hypothesized that *C. albicans* strains could be differentially sensitive to CRAMP which may in turn affect their ability to colonize the GI niche. To test this hypothesis, 529L, CHN1 and SC5314 were grown both aerobically and anaerobically at 37°C with different concentrations of the CRAMP peptide and growth rates were monitored in real time (aerobic) or as endpoints (anaerobic). Under aerobic conditions, SC5314 growth was substantially inhibited by low concentrations of CRAMP (5 µM) and no growth was observed with 10 µM CRAMP. In contrast, both 529L and CHN1 were more resistant to CRAMP and showed some ability to grow in the presence of 10 µM of this peptide (Figure 5A).

**Figure 5.**
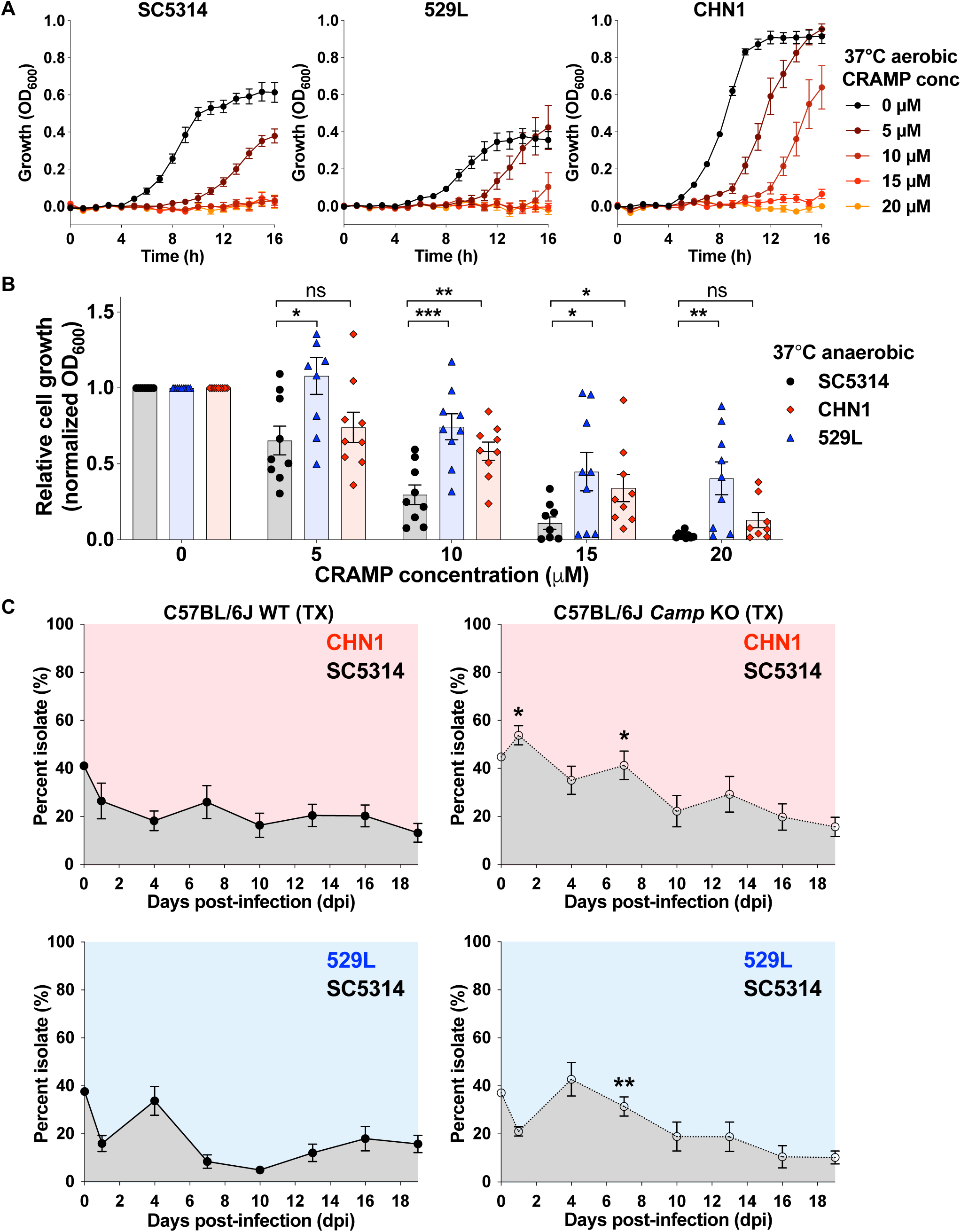
Effect of CRAMP on *C. albicans* growth and GI colonization. (A) *In vitro* susceptibility of *C. albicans* isolates SC5314, CHN1 and 529L to different CRAMP concentrations under aerobic growth at 37°C. Plots show means ± SEM growth levels over 16 h from 3 biological replicates. (B) *In vitro* susceptibility of *C. albicans* isolates to different CRAMP concentrations at 37°C under anaerobic conditions. Histograms show mean relative fungal growth ± SEM values from 3 biological replicates. * *P* < 0.05, ** *P* < 0.01, *** *P* < 0.001 based on comparison between SC5314 and CHN1 or 529L using unpaired parametric t tests. (C) Direct competitions between SC5314 and 529L or CHN1 in the GI of C57BL/6J wild type (WT) and *Camp* KO mice (both in TX). Isolates were co- inoculated in a 1:1 ratio and their proportions were determined using nourseothricin selection upon recovery from fecal pellets. Plots show mean values ± SEM from 8 mice per group, * *P* < 0.05, ** *P* < 0.01 based on comparisons of individual time points between WT and *Camp* KO mice using Mann-Whitney tests.

Similar trends were obtained when strains were grown anaerobically; SC5314 growth was reduced by ∼70% with 10 µM CRAMP whereas CHN1 and 529L showed a ∼42% and a ∼25% reduction in growth at this concentration (Figure 5B). Differences between strains were also observed at higher concentrations, with 529L being the most resistant to CRAMP (Figure 5B). This data establishes that SC5314 is significantly more sensitive to CRAMP than CHN1 and 529L under both aerobic and anaerobic conditions *in vitro*. Inspection of *XOG1*, the *C. albicans* gene which encodes for the β-(1, 3)-exoglucanase targeted by LL-37/CRAMP (27, 60), did not reveal genetic differences that could explain the differential sensitivity of the three isolates to this antimicrobial peptide (Supplemental Figure 7E).

To determine whether differences in CRAMP sensitivity could affect GI colonization, we performed direct competition experiments between SC5314 and CHN1 or SC5414 and 529L in both wild type C57BL/6J mice as well as in *Camp* knockout mice that lack the gene encoding the CRAMP peptide. Mice were gavaged with an equal mix of strains and the relative proportions of each strain determined by analyzing nourseothricin resistant/sensitive CFUs in fecal pellets every two days. Notably, we found SC5314 showed a relative fitness defect to CHN1 and 529L in mice regardless of whether they contained the *Camp* gene (Figure 5C). However, SC5314 was present at a significantly higher proportion of the population in *Camp* KO mice than in control mice at specific time points. Although a modest phenotype, this result implicates differences in GI colonization between CHN1/529L strains and SC5314 as being due, at least in part, to their differential susceptibility to the CRAMP antimicrobial peptide. This result is not unexpected given the variety of factors that promote *C. albicans* colonization resistance in the gut, including other antimicrobial peptides (e.g., β-defensins (61)), metabolites (e.g., short chain fatty acids (49)), and humoral factors (e.g., IgA (62)). As such, resistance to a single immune effector such as CRAMP would not be sufficient to completely explain the observed phenotypes.

## Discussion

*C. albicans* is a prevalent commensal of the human GI tract and yet is absent from the GI tract of most laboratory mouse strains. Moreover, colonization has typically required that adult mice are pretreated with antibiotics to enable stable colonization with SC5314, the standard *C. albicans* ‘laboratory’ isolate. Here, we demonstrate that two alternative clinical isolates, CHN1 and 529L, allow for long-term colonization of the gut of adult mice even without antibiotic supplementation, whereas SC5314 is gradually lost from the GI tract under the same conditions. Colonization is particularly stable in C57BL/6J mice which is the most widely used strain for biomedical research. We highlight that the increased stability of CHN1 and 529L over SC5314 was observed in multiple murine strains (C57BL/6J, BALB/c, C3H/HeN) and in three separate animal facilities located in New York, Rhode Island, and Texas. This establishes that the increased colonization fitness of CHN1/529L relative to SC5314 is a general finding that is not unique to a single animal facility or mouse line, and substantially expands the robustness of the current study. This finding suggests that CHN1 and 529L will also stably colonize the GI tract of mice in other animal facilities, establishing these strains as useful tools for researchers worldwide.

A range of murine models have been used to study *C. albicans* colonization, yet most of these models use sustained antibiotic treatment with adult mice which results in variable colonization levels (12, 19, 20, 63–65). Neonatal models that utilize infant mice (∼5-7 days of age) do not require antibiotic supplementation, which is attributed to an immature gut microbiota that lacks *Candida* colonization resistance (66, 67). Similarly, germ-free mice do not require antibiotics since they have no bacterial microbiota to inhibit *Candida* growth (21, 32, 68) . Finally, diet modification using a low fiber purified chow has also been shown to facilitate stable *Candida* gut colonization in mice even without antibiotics, presumably due to changes in the bacterial microbiome (18).

The current study highlights that intra-species variation has a major impact on *C. albicans* commensalism among other attributes. Several intra-species differences have previously been documented for *C. albican*s both *in vitro* as well as in systemic and oral infection models (36–38, 69). For example, while SC5314 is considered the standard lab isolate, this strain is one of the most virulent *C. albicans* strains in the murine systemic model (69) and shows a higher propensity to filament *in vitro* than other isolates (37). In most cases, the mechanisms by which intra-species variation impacts fungal cell behavior have not been defined, although decreased genome heterozygosity and homozygosity of the mating type-like (*MTL*) locus have both been linked with reduced systemic virulence (36, 70, 71).

Interestingly, the niche from which clinical *C. albicans* strains are isolated generally does not correlate with phenotypic properties, consistent with the idea that the same isolate can grow in multiple host tissues. Notable exceptions to this include a sub-clade of low heterozygosity strains (clade 13, *Candida africana*) that show decreased virulence in animal models of infection and may be restricted to genital tract infections (41, 72), and hyper-filamentous *nrg1* mutants that have been repeatedly recovered from the lungs of cystic fibrosis patients (73). Loss of filamentation ability has also been observed in some clinical isolates and can enhance GI colonization of antibiotic-treated mice (33, 59). Previous findings have therefore established that natural variation can impact *C. albicans*-host interactions, and the current study adds to this concept by identifying strains that show differences in GI fitness in the absence of antibiotic treatment.

We note that 529L was obtained from a patient with oral candidiasis (46) while CHN1 was isolated from the human lung (47), indicating that these strains were not isolated from the GI tracts of their respective hosts. Laboratory experiments have shown that 529L can persistently colonize the murine oral cavity, unlike SC5314 (46), and this was linked to a decreased inflammatory response to 529L (38). Additional studies have documented instances in which strain variation impacted immune responses to *C. albicans* during a systemic infection and highlighted differences in cell wall architecture as possible causes for strain-specific phenotypes (74).

The CHN1 isolate has not been studied extensively yet was previously shown to stably colonize the murine GI tract following pre-treatment with cefoperazone, a broad-spectrum antibiotic (47, 75). SC5314 and CHN1 colonization behavior was subsequently compared and both showed similar GI colonization properties in mice pre-treated with this antibiotic (47). The ability of these two strains to alter the bacterial microbiota following antibiotic treatment was also evaluated and both antagonized the re-growth of *Lactobacillus* (after cessation of antibiotic treatment) while promoting the growth of *Enterococcus*, indicating shared impacts on the bacterial microbiota (47). In the current study, we did not observe changes in the bacterial microbiota with colonization by SC5314, CHN1 or 529L. These differences in modulating the bacterial population are presumably due to differences in experimental design, with the current study showing that *C. albicans* colonization is not correlated with substantial changes to the composition of the bacterial microbiome.

Analysis of the *in vitro* phenotypes of SC5314, CHN1, and 529L revealed stark differences, with both CHN1 and 529L showing reduced metabolic and filamentation abilities relative to SC5314. 529L showed a particularly marked defect in growth and filamentation under a wide variety of conditions. However, all three strains showed similar propensities to filament in the GI niche, and 529L and SC5314 were previously shown to also exhibit similar filamentation phenotypes in the oral infection model (38). Our results indicate that *in vivo* filamentation characteristics can be very different from those observed *in vitro* and extend previous studies in which mutant *C. albicans* strains were shown to adopt different morphologies in the GI tract than those predicted based on *in vitro* phenotypes (35).

Sequencing of the 529L and CHN1 isolates did not reveal any obvious genetic alterations that might enable these strains to colonize mice better than SC5314. Thus, aneuploid configurations previously associated with increased fitness in the GI tract were not detected in these strains, although some homozygous tracts were shared by CHN1 and 529L that were absent in SC5314. However, the very large number of genetic differences between the three isolates makes identification of causal genetic links hard to establish without an extensive investigation of these differences.

It is likely that multiple mechanisms contribute to the observed strain differences in GI tract colonization. Genetic and phenotypic differences described here are likely to play important roles. We report one mechanism by which strain-specific differences in susceptibility to an intestinal-derived antimicrobial peptide (CRAMP) likely contribute to differences in colonization capacity, with SC5314 being more sensitive to this peptide than 529L/CHN1. Interestingly, certain prominent gut commensal bacteria (including Bacteroidetes) are also more resistant to gut-derived antimicrobial peptides when compared to gut pathobionts (e.g., *E. coli*), which can promote the dominance of commensal gut microbiota over pathobionts in the gut (76). Thus, the multiple factors (*e.g.*, genetic, phenotypic, environmental) that modulate *Candida* strain-specific differences in antimicrobial peptide sensitivity merit further investigation.

## Acknowledgements

We thank Mairi C Noverr for the gift of the CHN1 strain. This work was supported by National Institutes of Health (NIH) grants R01 AI093808 (TMH), R01 AI139632 (TMH), R21 AI105617 (TMH), R21 AI156157 (TMH), Burroughs Wellcome Fund Investigator in the Pathogenesis of Infectious Diseases Award (TMH), Ludwig Center for Cancer Immunotherapy (TMH), NIH P30 CA008748 (to MSKCC), NIH R01 AI123163 (AYK), K24 AI150992 (AYK), Roberta I. and Normal L. Pollock Fund (AYK), NIH R01 AI41893 (RJB), NIH R01 AI081704 (RJB), NIH R21 AI144651 (RJB), NIH R21 AI139592 (IVE), NIH NIGMS IDeA award P20GM109035 (IVE), Institut Pasteur G5 (IVE), CIFAR Azrieli Global Scholar Award (IVE), NIH NCCIH R21AT010366 (PB), NIDDK R01 DK125382 (PB) and NSF Graduate Research Fellowship award 1644760 (SP). The funding agencies had no role in the design and preparation of this manuscript.

## Author contributions

Designed research: LM AAM NT BZ RJB AYK TMH IVE; performed research: LM AAM NT PK SP SHL BZ IVE; analyzed data: LM AAM NT PK SP SHL CM BZ YT PB RJB AYK TMH IVE; wrote the paper: LM AAM NT RJB AYK TMH IVE.

## METHODS

### Growth of *C. albicans* isolates

All *C. albicans* isolates used in this study are listed in Supplemental Table 1. Unless otherwise specified, isolates were cultured overnight in 2-3 mL of liquid YPD (2% bacto-peptone, 1% yeast extract, 2% dextrose) at 30°C with shaking (200-250 RPM). Cell densities were measured using optical densities of culture dilutions (OD_600_) in sterile water using a Biotek Epoch 2 plate reader.

### Strain construction

To generate *SAT1*+ strains for GI competition assays, plasmid pDis3 was introduced into the *NEUT5L* neutral locus in the genome (48). The plasmid was linearized with NgoMIV and transformed into SC5314, CHN1 and 529L strains to generate *SAT1*+ transformants (Supplemental Table 1), which were selected on YPD+NAT (nourseothricin at 200 µg/ml, Werner Bioagents). PCR with primers 3118 (CCCAGATGCGAAGTTAAGTGCGCAG) and 4926 (AAAAGGCCTGATAAGGAGAGATCCATTAAGAGCA) from (48) was used to check correct integration of the *SAT1* gene.

### CRAMP *in vitro* assays

*C. albicans* isolates were grown overnight in Synthetic Complete Medium (SC) at 30°C under aerobic conditions. Cells were inoculated in 3 ml of liquid SC at OD_600_ 0.25, grown at 30°C until OD_600_ of 1, harvested by centrifugation and washed twice with 10 mM Sodium Phosphate Buffer pH 7.4 (NaPB). Cells were then resuspended in 3 ml of NaPB. 10 µl of cell resuspension was added to 140 µl YPD media with or without the desired concentration of CRAMP (Anaspec, AS-61305) and incubated for 1 h at 37°C with shaking. 40 µl of each culture was then added to an individual well of 96-well plate containing 60 µl YPD with the respective concentration of CRAMP. The plate was then incubated in a plate reader (Biotek Synergy HT) at 37°C with orbital shaking for 16 h. Growth was assessed by taking OD_600_ readings every hour. Aerobic experiments were performed with 3 biological experiments (with 3 technical replicates per biological experiment). For anaerobic growth in the presence of CRAMP, the 96- well plate was incubated at 37°C in an anaerobic chamber without shaking. Growth was evaluated by measuring the final biomass (OD_600_) at the end of the 16 h incubation period. Anaerobic experiments were performed with 3 biological experiments (with 3 technical replicates per biological experiment).

### Filamentation assays

For filamentation, *C. albicans* cells were grown overnight in YPD, washed in PBS and resuspended in PBS at a concentration of 10^5^ cells/ml. 1 ml of cell suspension was added to 24-well plates containing different media and plates were incubated for 4 h at 37°C with shaking. Images of approximately 500-1000 cells were captured using an AxioVision Rel. 4.8 (Zeiss) microscope. Assays were performed with 3 biological replicates.

### Phenotype microarray plate assays

*C. albicans* isolates were grown in YPD medium and then resuspended in sterile water to an OD_600_ of 0.2. The cell suspension was diluted 1:48 into inoculating fluid (IFY-0) and 100 µL of the cell suspension was aliquoted into each well of Biolog PM1 and PM2 plates according to manufacturer’s instructions (Biolog Inc., Hayward, CA). The plates were grown at 37°C for 24 h on a shaking platform at 200 RPM either aerobically or anaerobically (using Thermo Fisher AnaeroPack Anaerobic Gas Generators in a sealed plastic bag). Following incubation, wells were scored for filamentation on a scale of 1 to 5 with representing the proportion of filamentous cells in the population (1: 0-20%; 2: 20-40%; 3: 40-60%; 4: 60-80%; 5: 80-100%). PM experiments were performed with biological duplicates with growth (OD_600_) and filamentation scores averaged across the two replicates. Correlation analyses between growth and filamentation were performed using a simple linear regression model in GraphPad Prism 9.

### Gastrointestinal colonization and competition experiments

#### Experiments in Rhode Island

For animal infections, 7–8-week-old female BALB/c (stock 028, Charles River Laboratories) or C57BL/6J (stock 000664 from Jackson Laboratory, room MP14) female mice were housed together with free access to food (standard rodent chow, LabDiet #5010, autoclaved) and water. After 4 days of acclimation in the animal facility, mice were orally gavaged with 10^8^ cells and fungal cells were isolated from fecal pellets every other day by plating for CFUs. Pellets were homogenized in a PBS solution supplemented with an antibiotic mixture (500 µg/mL penicillin, 500 µg/mL ampicillin, 250 µg/mL streptomycin, 225 µg/mL kanamycin, 125 µg/mL chloramphenicol, and 125 µg/mL doxycycline). At the end of the experiment, mice were sacrificed and the number of fungal cells in each of the GI organs (stomach, small intestine, cecum, and colon) were determined by plating multiple dilutions of organ homogenates. For competition experiments, *C. albicans* cells were grown overnight in YPD at 30°C, washed with sterile water and quantified. 10^8^cells (containing a 1:1 ratio of each competing strain) were orally gavaged into the mouse GI tract. For each competition, one strain was nourseothricin sensitive (*SAT1-*) and one strain was nourseothricin resistant (*SAT1+*). Fecal pellets were collected every other day for 14 days, after which mice were euthanized and GI organs collected for CFU determination. Abundance of each strain was quantified by plating the inoculum, organ and fecal pellets homogenates onto YPD and YPD supplemented with nourseothricin (200 µg/ml, Werner Bioagents).

#### Experiments in Texas

For GI colonization experiments with single strain infection, 6-8 weeks old C3H/HeN female mice were bought from Envigo (stock 040, C3H/HeNHsd). Mice were fed Teklad Global 16% Protein Rodent Diet chow (Teklad 2916, irradiated). Mouse cages were changed once weekly. *C. albicans* isolates SC5314, CHN1 and 529L were grown overnight in YPD at 30°C with shaking under aerobic conditions. Cells were harvested, washed twice with PBS, and resuspended in PBS at a concentration of 1 x 10^9^ CFU/ml. C3H/HeN female mice were gavaged with 200 µl of cell suspension containing a total of 2 x 10^8^ *Candida* cells. To determine fungal burdens, fecal pellets were collected every 7 days for 35 days, homogenized and plated on YPD agar supplemented with antibiotics (30 µg/ml of vancomycin and 30 µg/ml of gentamicin).

For competition experiments, *C. albicans* isolates SC5314 (containing the *SAT1* gene, *SAT1*+), CHN1 and 529L were grown overnight in YPD at 30°C with shaking under aerobic conditions. Cells were harvested, washed twice with PBS and resuspended in PBS at a concentration of 1 x 10^9^ CFU/ml. Equal cell numbers of SC5314 (*SAT1*+) and CHN1 or 529L were mixed together. 6-8 weeks old C57BL/6J (Jackson Laboratories, room RB12) or *Cramp* KO (Jackson Laboratories, stock 017799) female mice were gavaged with 200 µl of cell suspension containing a total of 2 x 10^8^ *Candida* cells. Equal strain ratios were confirmed by plating the initial inoculum. Fecal pellets were collected every two days for 19 days, homogenized and plated on YPD agar supplemented with nourseothricin (200 µg/ml) and antibiotics (30 µg/ml of vancomycin and 30 µg/ml of gentamicin).

#### Experiments in New York

C57BL/6J (stock 00664, Jackson Laboratory, room MP14) female mice were purchased in groups of 20 mice and redistributed between cages to normalize gut microbiome one week prior to use. Mice were fed Lab Diet 5053 (PicoLab Rodent Diet 20, Irradiated). Mouse cages were changed once weekly. For GI colonization, *Candida* strains were streaked on SAB agar from glycerol stock and grown overnight at 37°C. Colonies were collected into YPD media and grown for an additional 18 h at 30°C with shaking (250 RPM). Cells were then collected in water and densities were measured using a hemocytometer. Mice were gavaged with 0.2 mL liquid culture containing a total of 10^7^ cells per mouse. Fecal samples were collected prior to gavage and regularly over 48 days during colonization. Gut fungal burdens were determined by plating fecal pellet homogenates on SAB agar (BD Difco Sabouraud Dextrose Agar, BD 210930) plates supplemented with 10 µg/ml of Vancomycin (Hospira, NDC 0409-6510-01) and 100 µg/ml of Gentamicin (Gemini, 400108).

For GI competitions, *SAT1*- and *SAT1*+ *Candida* strains were grown as for GI colonization experiments. Mice were gavaged with 5 x 10^6^ cells of each *SAT1*- and *SAT1*+ strains (total of 10^7^ cells per mouse). Fecal samples were collected regularly over 2-6 weeks and plated onto SAB and SAB with nourseothricin (100 μg/mL, Gold Biotechnology, N-500-1) plates.

### Analysis of *C. albicans* morphology in the mouse gut

*Candida* cells in the different GI sections were imaged by Fluorescence In Situ Hybridization (FISH) as described by (35). In brief, C57BL/6J mice were treated with an antibiotic cocktail (penicillin 1.5 mg/mL, streptomycin 2 mg/mL, 2.5% glucose for taste) and fluconazole (0.5 mg/ml, Sigma-Aldrich) for 3 days and followed by antibiotic treatment for one day. At this point, the mice were colonized by adding *C. albicans* cells (2 x 10^5^ cells/ml) to the drinking water containing antibiotics. The antibiotic containing water was changed every 3-4 days. After 7 days of colonization, the mice were sacrificed, and the GI organs were harvested. 1-2 cm pieces of different parts of the GI tract were fixed in methacarn (American Master Tech Scientific) overnight followed by two washes with 70% ethanol and subjected to paraffin block preparation. 10 μm sections were first deparaffinized and then the protocol from (35) was followed. *Candida* cells were stained with a Cy3-labelled PAN fungal 28s rRNA probe, epithelial cells were stained with DAPI (Molecular Probes, Invitrogen), and the GI mucosal layer was stained with Fluorescein labelled WGA1 and UEA1 (Vector Laboratories). Tissue imaging was carried out using colon sections and images were captured using an AxioVision Rel. 4.8 (Zeiss) fluorescence microscope. 8-10 Z-stacks were merged to generate the final images.

To evaluate *Candida* morphology in the GI, 10 μm tissue sections were first deparaffinized, blocked with 1X PBS + 5% FBS for 30 min at room temperature and then incubated with an anti-*Candida* antibody coupled to FITC (1:500 dilution, BIODESIGN International) overnight at 4°C. This was followed by 3 washes with PBS at room temperature and then staining of the epithelium with DAPI. Cell counting was carried out using an AxioVision Rel. 4.8 (Zeiss) fluorescence microscope. Two tissue sections from each mouse (n = 3 mice) were imaged and 50-600 *Candida* cells per mouse were examined for morphology. The proportions of yeast and hyphal morphotypes were averaged for the two sections for each segment of the GI tract.

### Whole genome sequencing

To extract genomic DNA, isolates were grown overnight in YPD at 30°C and DNA isolated from ∼10^9^ cells using a Qiagen Genomic Buffer Set and a Qiagen Genomic-tip 100/G according to manufacturer’s instructions. Libraries were prepared using the Nextera XT DNA Library preparation kit protocol (Illumina) with an input of 2 ng/μL in 10 μL. Each isolate was sequenced using Illumina HiSeq 2000 generating 101 bp paired reads. The nuclear genome sequences and General Feature Files (GFF) for *C. albicans* SC5314 reference genome (version A22) were downloaded from the *Candida* Genome Database (http://www.candidagenome.org/). Alignment, coverage, ploidy, heterozygosity and variant calling were performed as previously described (77). Average coverage levels for SC5314, 529L and CHN1 were 141X, 466X and 245X, respectively. Heterozygosity plots were constructed using methods from (41). Phylogenetic assignment was performed using RaxML (78) as described by (41) and using the isolates from the same study to classify the strains. Large homozygous tracts were confirmed by visual inspection in IGV (79). Mutations in *XOG1* were identified using GATK4 (80) and manually inspected in IGV. Genetic variants identified between SC5314 versus 529L/CHN1 are included in Supplemental Tables 3 and 4.

### 16S Sequencing

#### Experiments in Rhode Island

DNA was extracted from samples using the ZymoBIOMICS Fecal/Soil DNA 96 Kit from Zymo Research (D6011, Irvine, CA) as per the manufacturer instructions. Total DNA was eluted in nuclease-free water and quantified using the dsDNA-HS on a QubitTM 3.0 fluorometer (Thermo Fisher Scientific, Waltham, MA). The 16S rRNA V4 hypervariable region was amplified from DNA using the barcoded 515F forward primer and the 806Rb reverse primers from the Earth Microbiome Project (81). Amplicons were generated using 5X Phusion High-Fidelity DNA Polymerase under the following cycling conditions: initial denaturation at 98°C for 30 s, followed by 25 cycles of 98°C for 10 s, 57°C for 30 s, and 72°C for 30 s, then a final extension at 72°C for 5 min. Gel electrophoresis was used to confirm the amplicon size. The pooled amplicon library was sequenced at the Rhode Island Genomics and Sequencing Center at the University of Rhode Island (Kingston, RI) on the Illumina MiSeq platform with paired-end sequencing (2 x 250 bp), using the 600-cycle kit. Raw 16S rRNA reads were subjected to quality filtering, trimming, de-noising, and merging using the Qiime2 pipeline (version 2018.11) (82). Taxonomic classification was done using the pre-trained Naive Bayes classifier and the q2-feature-classifier plugin trained on the SILVA 132 99% database. Beta diversity was calculated using the phyloseq package (version 1.30.0) (83) in R (version 3.6.2) and visualized using PCoA with a Bray-Curtis test. Raw sequence data were uploaded and made available on the NCBI Sequence Read Archive under BioProject number PRJNA735873.

#### Experiments in New York

16S DNA was extracted and purified from fecal samples collected days 0 (before *Candida* gavage), 5, 12, and 48 post *Candida* gavage with a QIAamp kit (catalog no. 51306). The V4/V5 16S rDNA region was then PCR-amplified using modified universal bacterial primers. PCR products were sent to IGO (Integrated Genomics Operation) for Illumina sequencing and library preparation. The sequences were then compared to the NCBI RefSeq RNA library and raw reads were preprocessed using DADA2 implemented in R. DADA2 was used to perform quality filtering on resulting sequences, infer exact amplicon sequence variants (ASVs) resulting sequences, and to filter and remove chimeras (84). A minority of samples of insufficient quality were excluded from the analysis. Taxonomic assignment to species level was performed using an algorithm incorporating nucleotide BLAST (85), with NCBI RefSeq (86) as reference training set. The ASV tables, taxonomic assignment, and sample metadata were assembled using the phyloseq package construct (83). Construction of the sequence table and phyloseq object, and all subsequent end-analyses were performed using R (version 3.4). Raw sequence data were uploaded on the NCBI Sequence Read Archive under BioProject number PRJNA734639 (see Supplemental Table 2 for associated metadata). Beta diversity was visualized using PCoA with a Bray-Curtis test. Between-group differences were tested using a permanova (Adonis function via the Vegan package in RStudio 1.4) (87).

### Data availability

Strains and plasmids are available upon request. Whole genome sequencing data for SC5314, 529L, CHN2 are available at NCBI SRA as BioProject PRJNA730828. The raw sequence reads for SC5314 and 529L have been previously published on NCBI under BioProject PRJNA193498 (37) for SC5314, and under accession numbers SRX276261 and SRX276262 for 529L (57). 16S raw reads are available on NCBI under BioProject numbers PRJNA734639 and PRJNA735873.

## Supplemental Figure and Table Legends

**Supplemental Figure 1.**
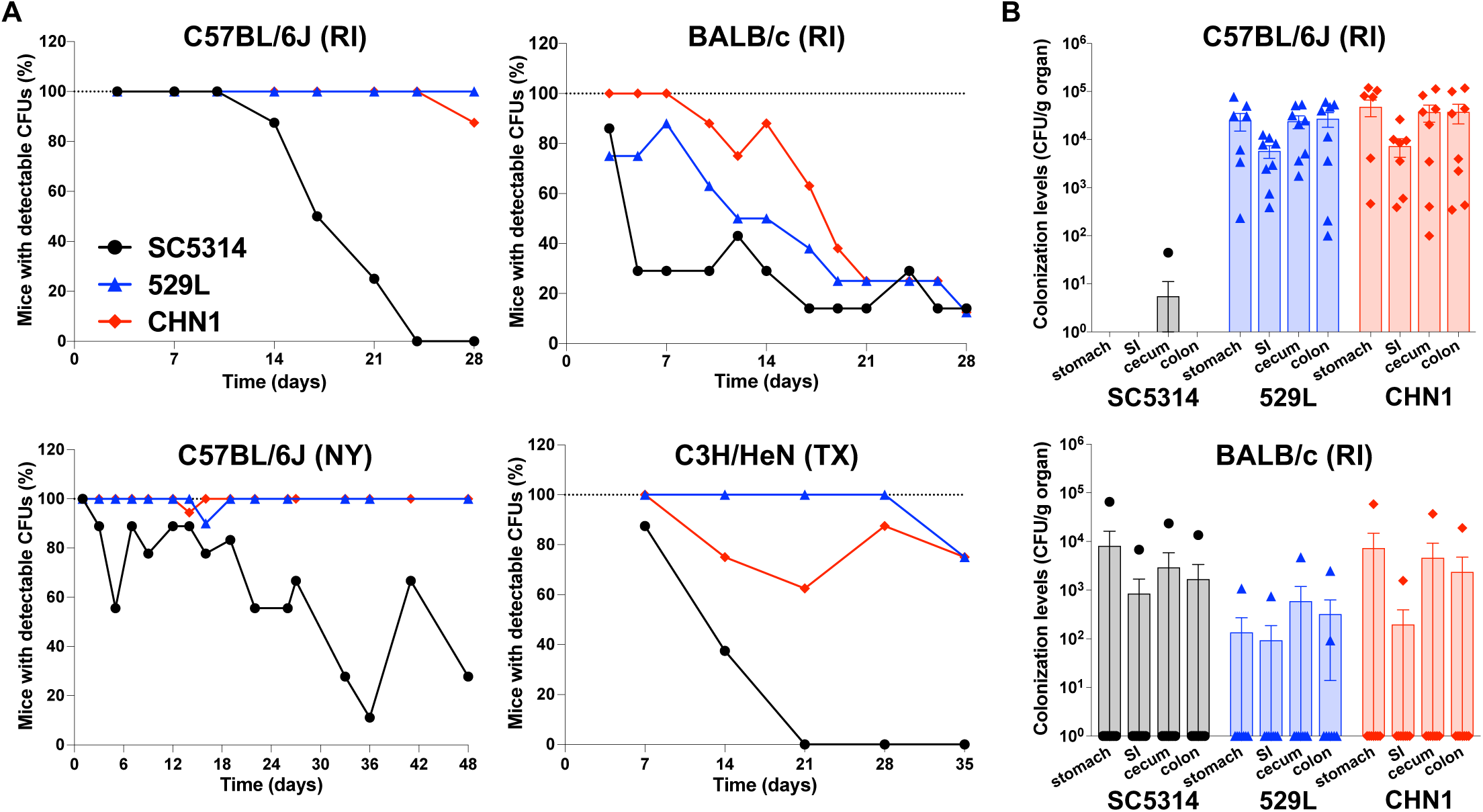
(A) Percent of mice with detectable fecal *C. albicans* CFUs during GI colonization of C57BL/6J (RI and NY), BALB/c (RI) and C3H/HeN (TX) mice from Figure 1. (B) GI organ colonization levels by isolates SC5314, 529L, CHN1 at the end of colonization (day 28) in C57BL/6J and BALB/c mice (RI) from (A). Panels show means ± SEM.

**Supplemental Figure 2.**
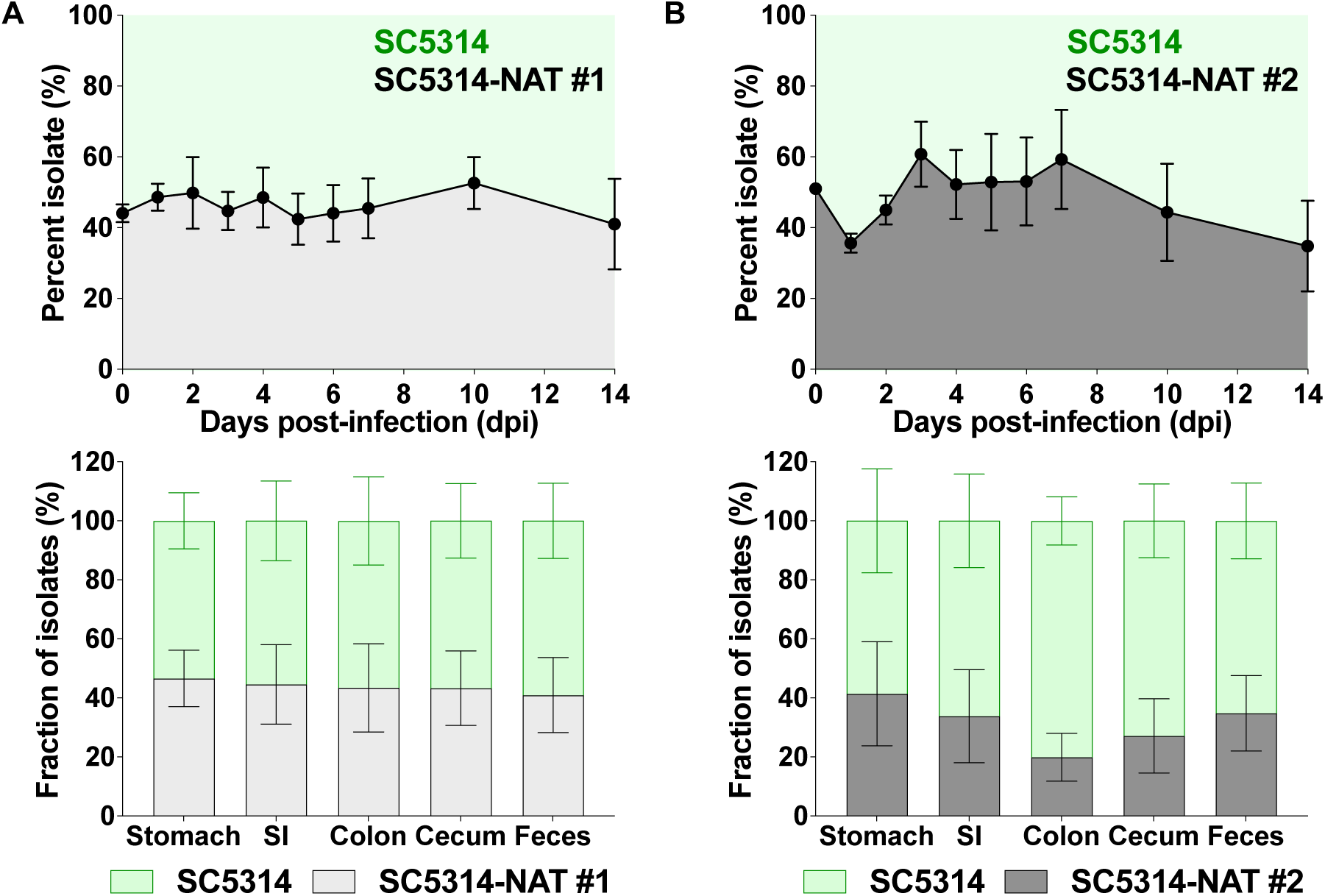
Integration of *SAT1* at the *NEUT5L* locus does not affect fitness in the mouse GI. No significant differences were observed when SC5314 and SC5314-*SAT1+* were directly competed in the GI tract of BALB/c mice (RI). Two independently transformed SC5314-*SAT1*+ strains (#1, #2) were used for these experiments (A, B). Strains were gavaged in 1:1 ratios and colonization levels were monitored for 14 days. After 14 days, the proportion of each strain was quantified from fecal pellets the GI tract organs. Histograms show means ± SEM from 7 single housed mice for each strain mix.

**Supplemental Figure 3.**
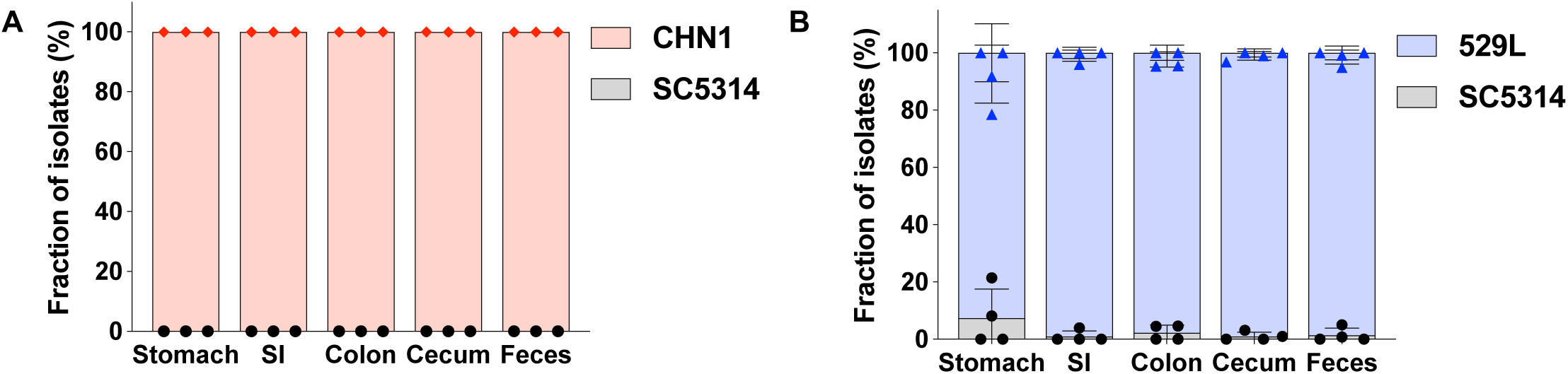
CHN1 (A) and 529L (B) outcompete SC5314 (*SAT1+*) in the GI organs of C57BL/6J mice (RI). Strains were gavaged in 1:1 ratios and colonization levels were monitored for 14 days. After 14 days, the proportion of each strain was quantified from the GI organs. Histograms show means ± SEM from 4 single housed mice.

**Supplemental Figure 4.**
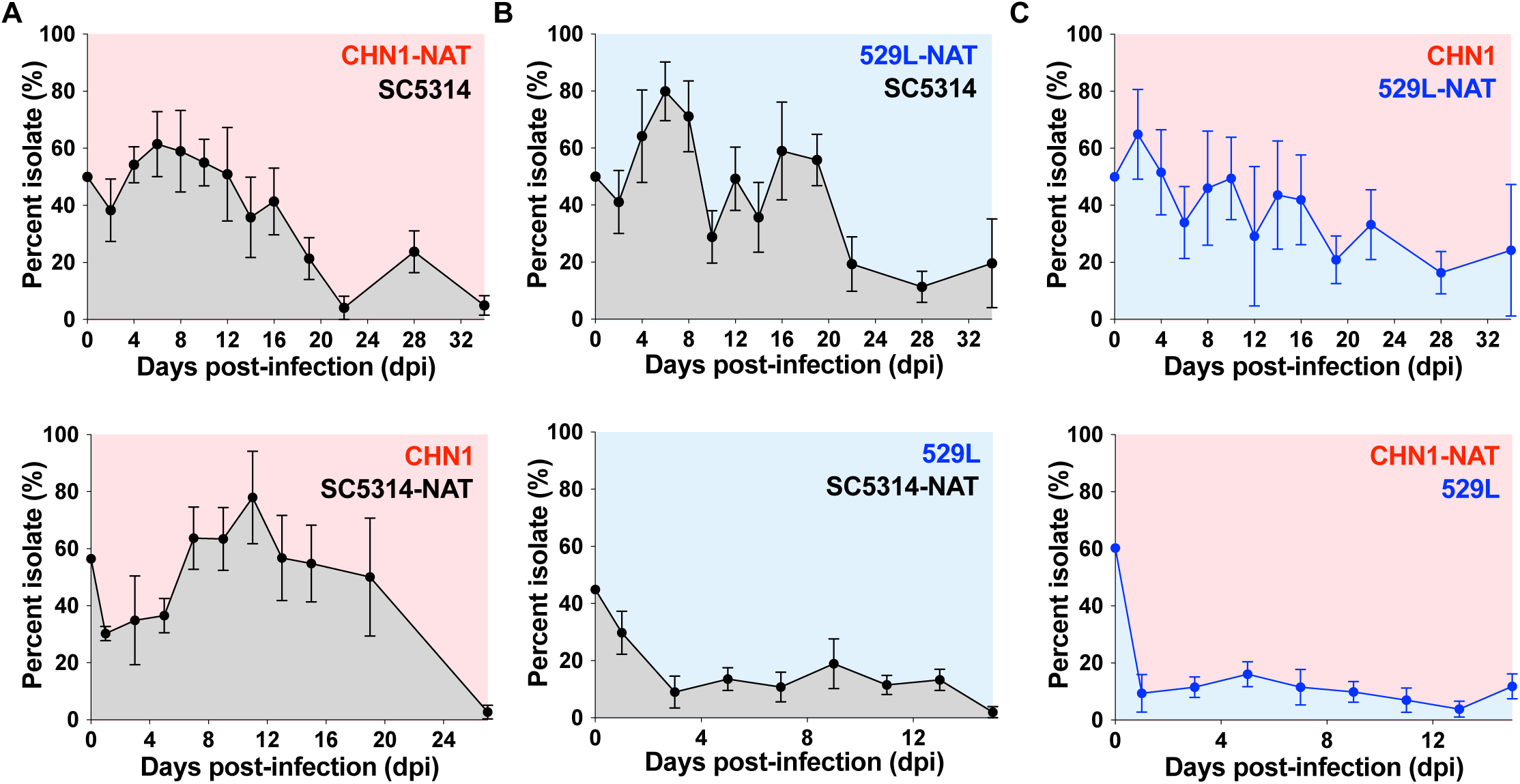
(A-B) CHN1 and 529L outcompete SC5314 in the GI tract of C57BL/6J mice (NY). Strains were gavaged in 1:1 ratios and colonization levels were monitored for 15-34 days. The proportion of each strain (%) was quantified from fecal pellets every 2-7 days using nourseothricin selection. (C) Direct competitions between CHN1 and 529L in the GI of C57BL/6J mice (NY) using the same method. Plots show means ± SEM from 5 mice.

**Supplemental Figure 5.**
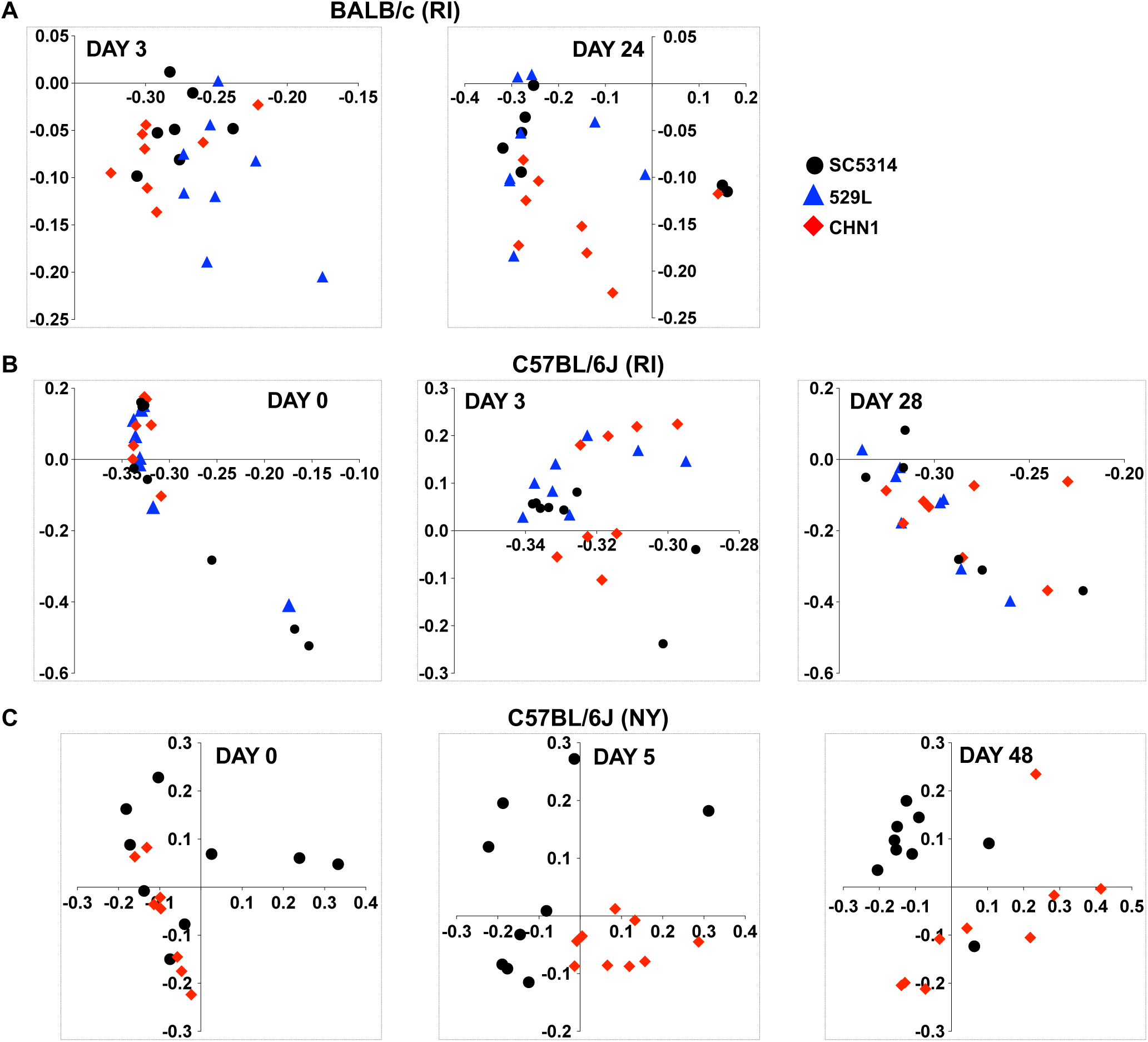
Bray Curtis PCoA plots showing strain effects for BalbC (A, RI) and C57BL/6J (B, RI; C, NY) mice colonized with strains SC5314, 529L and CHN1 prior to gavage and during GI colonization. No significant clustering of mice colonized with the same isolate was identified.

**Supplemental Figure 6.**
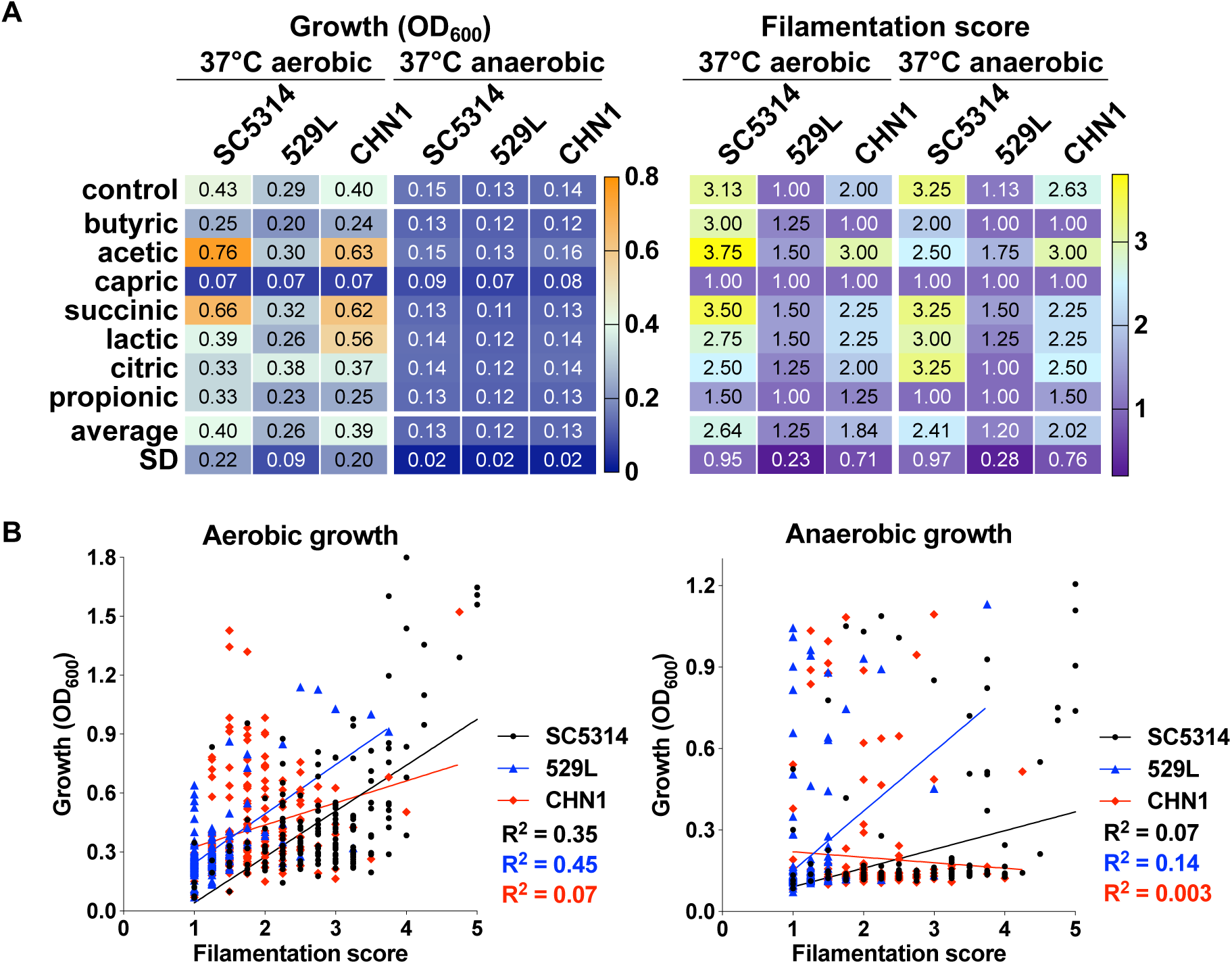
(A) Growth and filamentation of *C. albicans* isolates SC5314, CHN1 and 529L on 7 short chain carboxylic acids contained on Biolog PM plates. Heatmaps include control wells (no carbon source), as well as means ± SD values for each condition. (B) Correlation analyses between growth and filamentation under aerobic and anaerobic conditions for the three isolates. For each strain, R^2^ values represent the coefficient of determination indicating the goodness of fit for simple linear regressions.

**Supplemental Figure 7.**
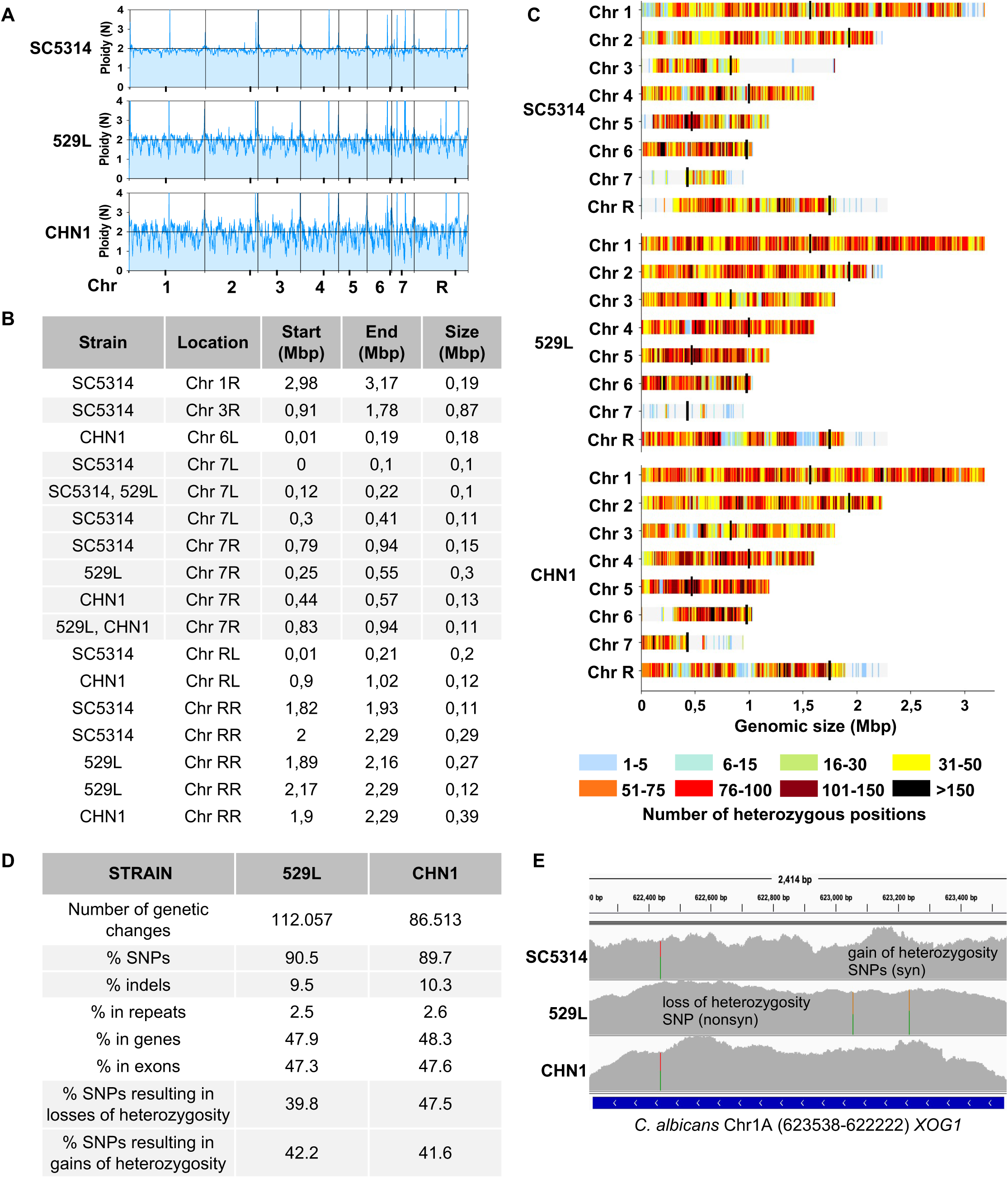
Genome sequencing of *C. albicans* SC5314, 529L and CHN1 illustrates extensive genetic differences between isolates. (A) Approximate ploidy levels for strains SC5314, 529L and CHN1 across the 8 *C. albicans* chromosomes; black dots on the X axis indicate centromere positions. (B) Size and position of large homozygous tracts (>0.1 Mbp) identified in the three isolates relative to the SC5314 reference strain (assembly 22). L, R indicate the left and right chromosome arms, respectively. (C) Density maps of heterozygous positions for the three isolates, shown for each chromosome across 10 kbp windows. Black bars indicate centromere positions. (D) Number of genetic changes identified in 529L and CHN1 relative to the SC5314 version examined in this study. (E) Genetic changes identified in the *XOG1* gene in the three isolates relative to the SC5314 reference genome. Image shows IGV coverage tracts with positions different from the reference genome highlighted in color. The three sites reflect positions which differ in 529L relative to the other 2 isolates; syn, synonymous mutation; nonsyn – nonsynonymous mutation.

**Supplemental Figure 8.**
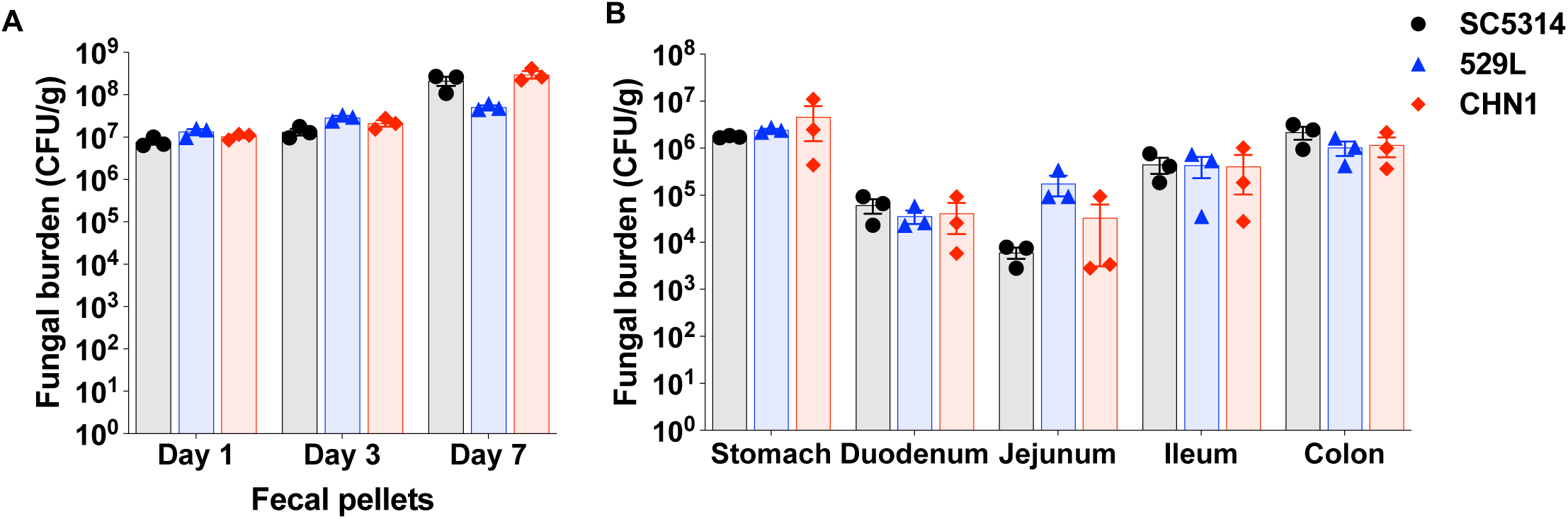
Fecal (A) and organ (day 7, B) fungal burdens of C57B/6J mice (RI) using an antibiotic model of colonization. Plots show means ± SEM from 3 single-housed mice.

**Supplemental Table 1.** Strains used in this study.

**Supplemental Table 2.** Metadata associated with NCBI BioProject PRJNA734639.

**Supplemental Table 3.** Genetic variants identified in isolate 529L relative to SC5314. For each variant, the table includes the type of mutation, genomic position and distances from the nearest gene, exon or repeat.

**Supplemental Table 4.** Genetic variants identified in isolate CHN1 relative to SC5314. For each variant, the table includes the type of mutation, genomic position and distances from the nearest gene, exon or repeat.

